# *ksrates*: positioning whole-genome duplications relative to speciation events in *K*_S_ distributions

**DOI:** 10.1101/2021.02.28.433234

**Authors:** Cecilia Sensalari, Steven Maere, Rolf Lohaus

## Abstract

**Summary:** We present *ksrates*, a user-friendly command-line tool to position ancient whole-genome duplication (WGD) events with respect to speciation events in a phylogeny by comparing paralog and ortholog *K*_S_ distributions derived from genomic or transcriptomic sequences, while adjusting for substitution rate differences among the lineages involved.

**Availability and implementation:** *ksrates* is implemented in Python 3 and as a Nextflow pipeline. The source code, Singularity and Docker containers, documentation and tutorial are available via https://github.com/VIB-PSB/ksrates.

## 1 Introduction

The *K*_S_ value of a pair of homologous sequences, i.e. the estimated number of synonymous substitutions separating them per synonymous site, is used as a proxy for the time elapsed since the sequences diverged (Lynch and Conery, 2000). *K*_S_ values of paralog pairs can be used to construct a relative age distribution of duplication events in a species, offering insight into the species’ gene duplication history. Distinctive peaks in such paralog *K*_S_ distributions are often used to infer the presence of whole-genome duplications (WGDs) (Blanc and Wolfe, 2004). Ortholog *K*_S_ distributions on the other hand are informative of the age of divergence between two species. A common practice to assess the temporal order of speciation and duplication events, e.g. to characterize the phylogenetic position of inferred WGDs, is to superimpose ortholog and paralog *K*_S_ distributions in mixed plots (Blanc and Wolfe, 2004; Zhang *et al*., 2017; Li *et al*., 2018).

However, the sequence of events inferred from such naive mixed *K*_S_ plots is not always reliable. *K*_S_ estimates for events of the same absolute age may differ depending on the synonymous substitution rates in the lineages involved. In general, the paralog *K*_S_ distribution of a species of interest is therefore not directly comparable to distributions of ortholog *K*_S_ values involving other species, and their straightforward superimposition in a mixed plot can be misleading.

*ksrates* generates adjusted mixed plots of *K*_S_ distributions by rescaling ortholog *K*_S_ estimates of species divergence times to the paralog *K*_S_ scale of a focal species, producing shifts in the estimated *K*_S_ position of each speciation event proportional to the estimated substitution rate difference between the diverged lineage and that of the focal species. A use case is presented to show that the phylogenetic placement of hypothesized WGDs can be inferred more accurately from such adjusted mixed plots than from naive mixed plots.

## 2 Methods

### 2.1 *K*_S_ distribution construction and mixture modeling

*ksrates* uses the *wgd* package (Zwaenepoel *et al*., 2019) to detect paralog pairs and one-to-one ortholog pairs from genomic or transcriptomic sequence data and to calculate the associated *K*_S_ values, and then constructs *K*_S_ distributions from the raw *K*_S_ data (see Supplementary Methods). When genome structural annotation (GFF3 file) is provided as input for the focal species, the i-ADHoRe 3.0 package (Proost *et al*., 2012) is run to identify anchor pairs, i.e. pairs of paralogs located on collinear duplicated segments remaining from WGDs. *ksrates* then uses mixture modeling on the anchor-pair *K*_S_ distribution or the whole-paranome *K*_S_ distribution of the focal species to detect potential WGD signatures. When anchor-pair data is available, the anchor-pair *K*_S_ values are associated with putative WGDs based on lognormal mixture model clustering of the median *K*_S_ values of collinear segment pairs (see Supplementary Methods and Fig. 1A). Otherwise, an exponential-lognormal mixture model is fit to the whole-paranome *K*_S_ distribution, with an exponential component capturing the small-scale duplication background and lognormal components modeling putative WGD peaks. Optionally, it is also possible to fit more simple lognormal-only mixture models to the whole-paranome and anchor-pair *K*_S_ distributions. The reader should be aware that mixture modeling results should be interpreted cautiously because mixture models tend to overestimate the number of components present in the target *K*_S_ distribution and hence the number of WGDs (Tiley *et al*., 2018).

**Figure 1.**
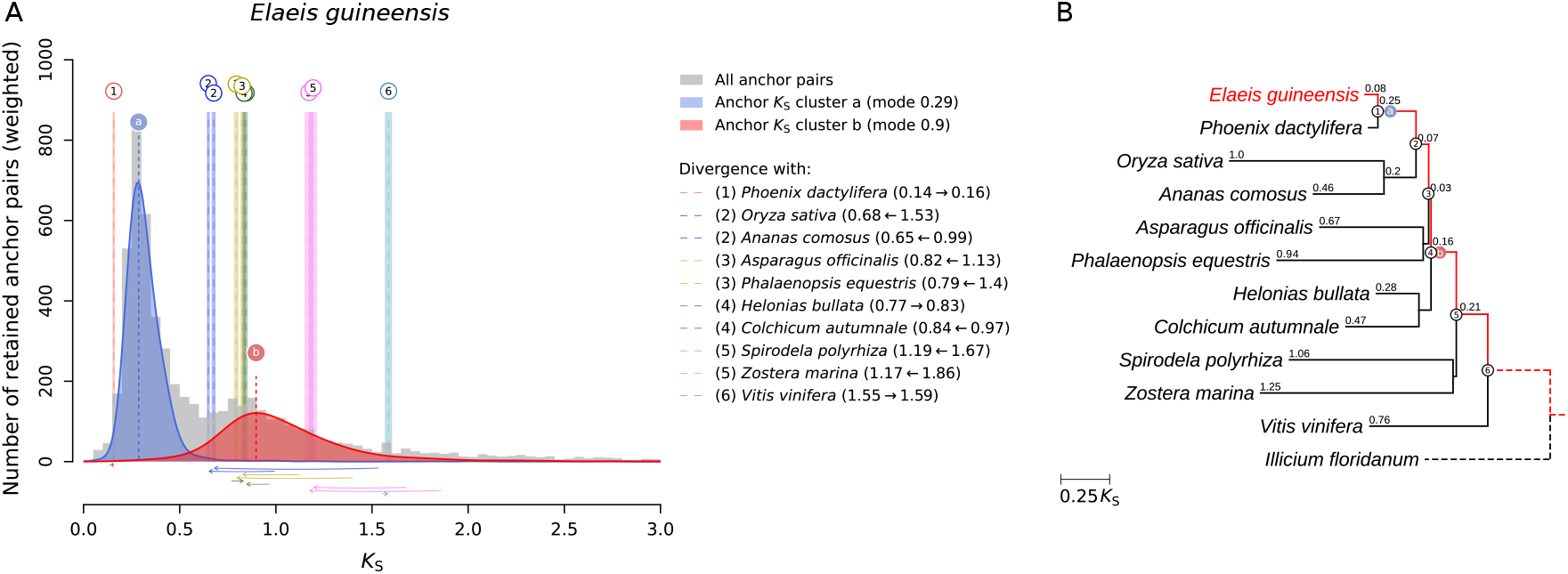
Use case analyzing WGD signatures in oil palm (*Elaeis guineensis*). (**A**) Substitution-rate-adjusted mixed paralog–ortholog *K*_S_ plot for oil palm as produced by *ksrates*. The anchor-pair *K*_S_ distribution for oil palm is shown in gray, with two putative WGD components inferred by lognormal mixture model clustering (see Supplementary Methods) indicated in blue and red. The vertical dashed lines labeled ‘a’ and ‘b’ indicate the modes of these components, which are taken as *K*_S_-based WGD age estimates. Rate-adjusted mode estimates of ortholog *K*_S_ distributions between oil palm and other species, representing speciation events, are drawn as numbered vertical long-dashed lines, with associated colored boxes ranging from one standard deviation (sd) below to one sd above the mean mode estimate. Lines representing the same speciation event in the phylogeny (numbered circles in (**B**)) share color and numbering. Horizontal arrows at the bottom of the plot indicate the speciation-line shifts produced by *ksrates’* substitution rate adjustments (compare to Supplementary Fig. 1), and the corresponding *K*_S_ value shifts are given in the panel legend. (**B**) Phylogram generated by *ksrates* from the input phylogenetic tree, with branch lengths set to the *K*_S_ distances estimated from ortholog *K*_S_ distributions. Short branches are only labeled for the *E. guineensis* lineage (highlighted in red). Numbered circles (added manually) indicate the speciation events along this lineage (numbering as in (**A**)). The blue and red filled circles labeled ‘a’ and ‘b’ (added manually) indicate the WGDs inferred in (**A**).

### 2.2 Ortholog *K*_S_ adjustment and output

Phylogenetic relationships between species are provided as an input phylogenetic tree. Raw *K*_S_ estimates for the divergence times between the focal species and other species are obtained by estimating the mode of the corresponding ortholog *K*_S_ distributions using bootstrapped kernel density estimation. For each divergence event, a substitution-rate-adjusted *K*_S_ estimate is then calculated by decomposing the raw *K*_S_ value into branch-specific contributions with the help of an outgroup species, similar to the methodology used in relative rate testing (Sarich and Wilson, 1973), and rescaling the contribution of the diverged species to the *K*_S_-timescale of the focal species (see Supplementary Methods). The adjusted divergence *K*_S_ estimates are then superimposed on the paralog *K*_S_ distribution of the focal species to produce an adjusted mixed *K*_S_ distribution plot (Fig. 1A).

## 3 Use case

Shown in Fig. 1A is a *ksrates* mixed plot of the anchor-pair paralog *K*_S_ distribution for the mono-cot plant *Elaeis guineensis* (oil palm) and substitution-rate-adjusted ortholog *K*_S_ peak estimates representing divergence events with other flowering plant species (see Fig. 1B). The two peaks detected in the anchor pair *K*_S_ distribution are caused by a known palm-specific WGD (Singh *et al*., 2013) (blue peak) and a much older WGD (*τ*, red peak) thought to be shared by all monocots except a few early-diverging clades such as the Alismatales (here represented by *Spirodela polyrhiza* and *Zostera marina*) (Ming *et al*., 2015). Because palms have a much lower substitution rate than other monocots such as grasses or Asparagales (Barrett *et al*., 2015, also Fig. 1B), a naive mixed paralog–ortholog *K*_S_ plot (Supplementary Fig. 1) however suggests that the older *τ* WGD would be a second palm-specific WGD event. Moreover, the naive mixed plot contains widely different *K*_S_ estimates for the same divergence event and produces an order of divergences inconsistent with the known phylogeny of flowering plants (Supplementary Fig. 1). Substitution rate adjustment with *ksrates* shifts the ortholog *K*_S_ estimates of most divergence events leftward towards younger *K*_S_ values, groups estimates of the same event more closely together and puts them in the commonly accepted order (Fig. 1B), resulting in a correct phylogenetic positioning of the two WGDs in the oil palm lineage.

## Data availability

The data underlying this article are available in Zenodo, at https://dx.doi.org/10.5281/zenodo.4717851.

## Author contributions

R.L. conceived the *K*_S_ rate-adjustment methodology. R.L. and S.M. designed tool functionality.

C.S. and R.L. implemented the tool. R.L. and S.M. supervised the study. All authors contributed to writing the manuscript.

## Acknowledgements

We thank Bert Droesbeke for technical assistance with the Singularity and Docker containers.

