## Supplementary Material for "*ksrates*: positioning whole-genome duplications relative to speciation events in *K*_S_ distributions"

* co-last authors

**3 Data sources** **14**

**4 Parameters used** **14**

**5 Software availability and requirements** **16**

**6 Supplementary references** **17**

### 1 Supplementary figures

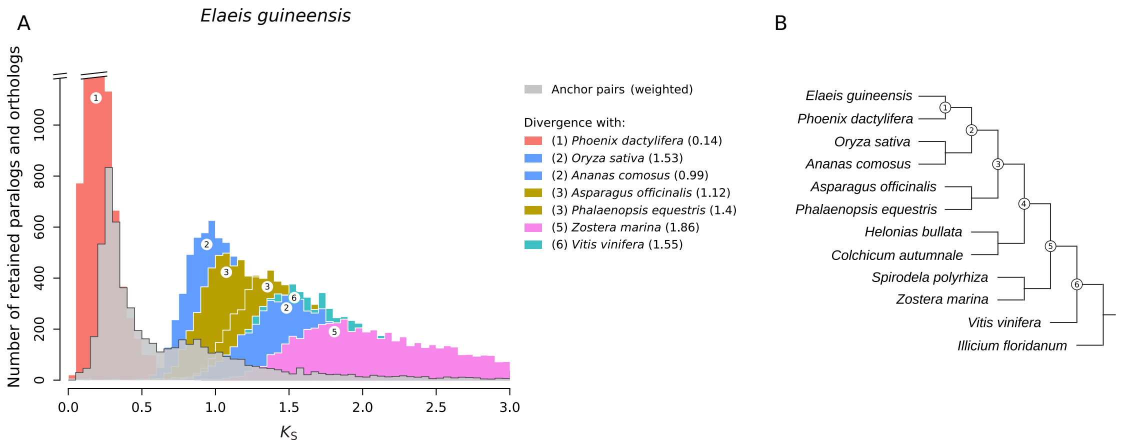

**Supplementary Figure 1.** Panel A shows a naive mixed paralog–ortholog $K_S$ plot for oil palm (*Elaeis guineensis*) without any substitution rate adjustment. The paralog $K_S$ distribution for oil palm anchor pairs is shown in gray. Ortholog $K_S$ distributions representing the same speciation event in the phylogeny (numbered circles in panel B) share color and numbering. Panel B shows the input phylogenetic tree. Numbered circles indicate the speciation events along the evolutionary lineage of oil palm (numbering as in A). The ortholog $K_S$ distributions for some species pairs in the input tree were omitted in A to improve visual clarity. Compare to Fig. 1 in the main text.

### 2 Supplementary methods

#### 2.1 Construction of paralog and ortholog $K_S$ distributions

The calculation of paralog and ortholog $K_S$ values is performed by the *wgd* package (Zwaenepoel *et al.*, 2019), which largely follows the approach outlined in (Vanneste *et al.*, 2013). Genomic or transcriptomic CDS sequence data for each species need to be provided in FASTA format. Briefly, one-to-one orthologs are recovered from all-versus-all BLASTp searches using the reciprocal best BLAST hit criterion. Paralogous gene families for the focal species are constructed from all-versus-all BLASTp results using MCL clustering (Enright *et al.*, 2002). Paralogous gene families with more than 200 members are excluded by default from further analysis (customizable parameter). For the multiple sequence alignment step, *ksrates* uses the default aligner in *wgd*, MUSCLE (Edgar, 2004). $K_S$ estimates of paralog and ortholog pairs are then calculated as described in (Zwaenepoel *et al.*, 2019). To compensate for the $K_S$ estimate redundancy in paralog gene families (multiple $K_S$ values may be estimated for the same duplication event (Maere *et al.*, 2005)), phylogenetic relationships in a gene family are reconstructed through FastTree (Morgan *et al.*, 2009), the default tool choice in *wgd*. Node weighting is then applied for constructing the paralog $K_S$ distribution(s) for the focal species as described in (Maere *et al.*, 2005), using weights derived from the phylogenetic relationships obtained by FastTree. *ksrates* generates $K_S$ distributions up to $K_S$=5 for paralogs and $K_S$=10 for orthologs by default (customizable parameters). If available, the genome structural information file (GFF3 file) for the focal species can also be provided to perform synteny/collinearity analysis and construct an anchor-pair $K_S$ distribution (Supplementary Fig. 2). Anchor pairs are a subset of paralog pairs found in duplicated genomic regions with conserved gene order (collinear segments), which likely originated through a whole-genome duplication (WGD)* or other large-scale duplication. i-ADHoRe (Proost *et al.*, 2012) is used to detect such collinear segments and their anchor pairs. $K_S$ values for these anchor pairs are extracted from the full set of $K_S$ values for all paralog pairs, reweighted, and assembled in an anchor-pair $K_S$ distribution.

*or, more generally, whole-genome multiplication (WGM), but we will here simply use the more commonly-used acronym WGD to refer to any multiplication

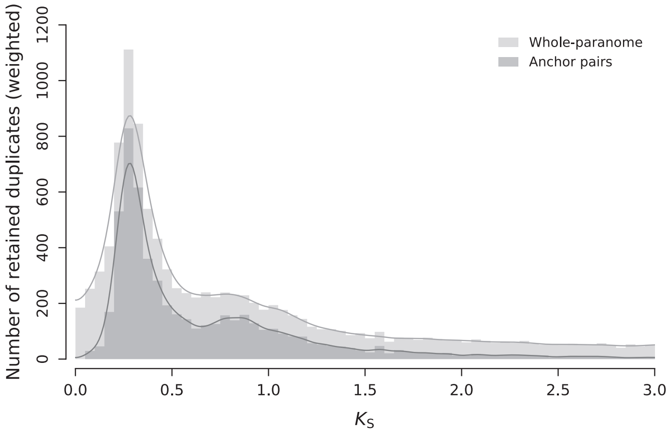

**Supplementary Figure 2.** Whole-paranome $K_S$ distribution (light gray histogram and KDE curve) and anchor-pair $K_S$ distribution (dark gray histogram and KDE curve) for oil palm (*Elaeis guineensis*). Two WGD peaks are visible on the left side of the distribution.

#### 2.2 Ortholog $K_S$ adjustment

The methodology for adjusting ortholog $K_S$ estimates is composed of several steps (Supplementary Fig. 3). *ksrates* first detects substitution rate differences between the focal species and the other species using ortholog $K_S$ estimate decomposition, and then subsequently rescales the ortholog $K_S$ estimates of species divergence events to adjust for such differences.

**Ortholog $K_S$ estimate decomposition** Ortholog $K_S$ adjustment requires known phylogenetic relationships between the focal species and all other species in the dataset. These relationships are provided to *ksrates* as input using the Newick tree format. Supplementary Fig. 3 shows an example input phylogenetic tree in which A is the focal species. To adjust the $K_S$ estimate of each divergence event between the focal species and other species in the phylogeny, each raw ortholog $K_S$ estimate is decomposed into branch-specific contributions. For example, the $K_S$ estimate of the divergence of focal species A and species B in the example tree is decomposed into the contributions from branches O–A and O–B, with O being the last common ancestor of species A and B. For this decomposition an outgroup species is needed (here, e.g. C) and hence each ortholog species pair to be included in the mixed paralog–ortholog $K_S$ plot is required to have at least one outgroup.

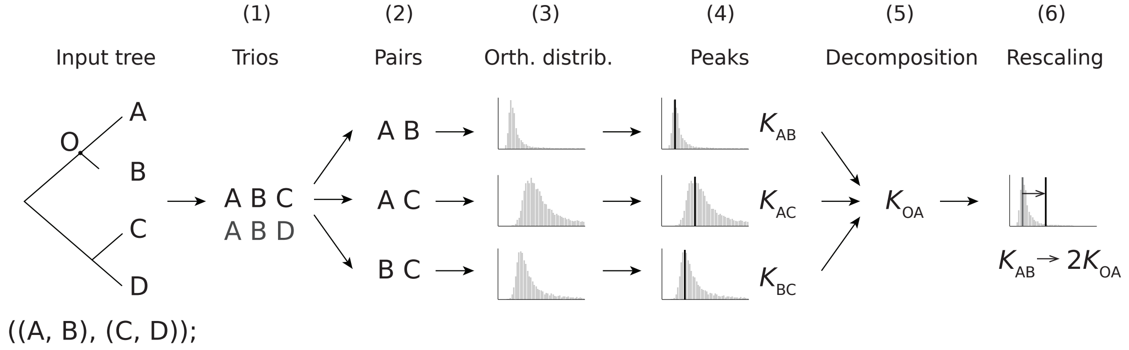

**Supplementary Figure 3.** The steps of the substitution-rate-adjustment methodology for ortholog $K_S$ estimates. Species A in the input tree is the focal species, O indicates the divergence event (or the common ancestor) of species A and B, and species C and D are outgroup species to the ingroup species pair A and B.

In the first step, the input phylogenetic tree is broken down into species trios, each composed of the focal species A, a diverged species in the dataset (here, B) and an outgroup reference (e.g. C). If the tree is large or complex enough it is possible to have more than one outgroup (here, e.g. D), and thus multiple trios, per diverged ingroup species pair. Each species trio is then broken down into its three possible species pairs (step 2), namely the pair of the two ingroup species (here, A and B), and two pairs consisting of each ingroup species together with the outgroup (here, e.g. A and C, and B and C). For each species pair of each species trio, the one-to-one orthologs are detected and their $K_S$ values estimated using the *wgd* package, and an ortholog $K_S$ distribution is built (step 3, see also Section 2.1 and Supplementary Fig. 4). A single ortholog $K_S$ estimate for the divergence time of each species pair is then obtained from its ortholog $K_S$ distribution (step 4). We use bootstrapped kernel density estimation (KDE) to estimate the mode of the ortholog $K_S$ distribution. The ortholog $K_S$ data is bootstrapped 200 times (customizable parameter) and each time the mode of a KDE with Gaussian kernel is computed. The final divergence $K_S$ estimate is then calculated as the mean of the bootstrapped modes together with an associated standard deviation (Supplementary Fig. 4). We denote the three ortholog $K_S$ estimates for the divergences between the three species pairs in our example trio as $K_{AB}$, $K_{AC}$ and $K_{BC}$. Note that $K_{AC}$ and $K_{BC}$ are $K_S$ estimates for the same divergence event in the evolutionary history of the three species (see example tree in Supplementary Fig. 3).

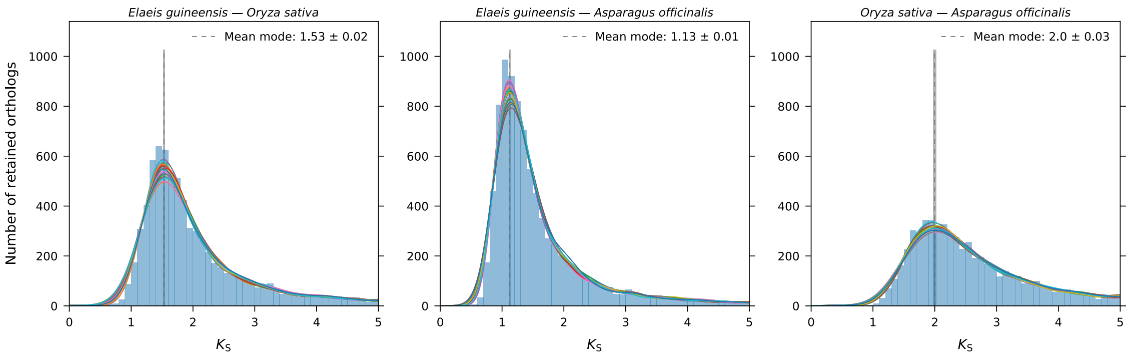

**Supplementary Figure 4.** The three ortholog $K_S$ distributions generated from the trio *Elaeis guineensis*, *Oryza sativa* and outgroup *Asparagus officinalis*. The ortholog $K_S$ histograms are shown in light blue, and the first 20 bootstrapped KDEs are depicted as colored curves. The estimated mean mode of each distribution, i.e. the ortholog $K_S$ estimate for the divergence time between the two species, is shown as a dashed gray line and its $K_S$ value is given in the legend. Colored boxes around these estimates range from one standard deviation (sd) below to one sd above the mean mode estimate. Note that the *Elaeis guineensis–Asparagus officinalis* $K_S$ distribution has a much younger mode than the *Oryza sativa–Asparagus officinalis* $K_S$ distribution, even though they represent the same divergence event, suggesting that the synonymous substitution rate in the lineage leading to *Elaeis guineensis* is lower than the rate in the lineage leading to *Oryza sativa*.

To decompose the ortholog $K_S$ estimate of the ingroup species pair, $K_{AB}$, we use simple equations from relative rate testing (Graur, 2016; Sarich and Wilson, 1973) (step 5). The ortholog $K_S$ estimate $K_{AB}$, i.e. an estimate of the average number of synonymous substitutions per synonymous site between the focal species A and a diverged species B, is the sum of the substitutions that occurred on branch O–A ($K_{OA}$) and on branch O–B ($K_{OB}$) (see example tree in Supplementary Fig. 3):

$$K_{AB} = K_{OA} + K_{OB} \tag{1}$$

With the help of the outgroup reference species in each trio (here, e.g. C), and the two corresponding ortholog $K_S$ estimates $K_{AC}$ and $K_{BC}$, we can calculate the value of $K_{OA}$:

$$K_{OA} = \frac{K_{AC} + K_{AB} - K_{BC}}{2} \tag{2}$$

$K_{\mathrm{OA}}$ is an estimate of the number of substitutions that occurred on branch O–A, i.e. the contribution of the branch O–A to $K_{\mathrm{AB}}$, and depends only on the synonymous substitution rate experienced in the lineage leading to focal species A. The error associated to $K_{\mathrm{OA}}$ can be derived from the estimated standard deviations ($s$) through error propagation rules:

$$s_{K_{\mathrm{OA}}} = \frac{\sqrt{s^2_{K_{\mathrm{AC}}} + s^2_{K_{\mathrm{AB}}} + s^2_{K_{\mathrm{BC}}}}}{2} \tag{3}$$

**Ortholog $K_{\mathrm{S}}$ estimate rescaling** To adjust the $K_{\mathrm{S}}$ estimate of the divergence of species A and B, $K_{\mathrm{AB}}$, to the $K_{\mathrm{S}}$-timescale of the focal species A we then simply rescale the contribution of the diverged branch $K_{\mathrm{OB}}$ to the same value as $K_{\mathrm{OA}}$, or, in short, we rescale $K_{\mathrm{AB}}$ to $2K_{\mathrm{OA}}$ (step 6):

$$K_{\mathrm{AB}} \rightarrow K_{\mathrm{OA}} + K_{\mathrm{OA}} = 2K_{\mathrm{OA}} \tag{4}$$

The error associated to the rescaled $K_{\mathrm{AB}}$ can be derived through error propagation rules:

$$s_{K_{\mathrm{AB}}} = \sqrt{s^2_{K_{\mathrm{OA}}} + s^2_{K_{\mathrm{OA}}}} = \sqrt{2}s_{K_{\mathrm{OA}}} \tag{5}$$

In other words, we calculate a rescaled $K_{\mathrm{AB}}$ from a hypothetical set of ortholog pairs that diverged at time point O and evolved under the same synonymous substitution rate history, the rate history experienced by the lineage leading to focal species A. Alternatively, to think of these adjustments of ortholog $K_{\mathrm{S}}$ estimates of species divergence times completely within the *paralog* $K_{\mathrm{S}}$-timescale of a focal species, one can imagine these pairs of sequences that diverge at a time point O not as ortholog pairs in diverging species but as paralog pairs that originate in a hypothetical duplication event in the lineage of the focal species A coinciding with the speciation of A and B.

When multiple ortholog $K_{\mathrm{S}}$ adjustments are computed for the same speciation event based on multiple outgroups, i.e. from species trios having the same two ingroup species but different outgroup species, the mean of all these rate-adjusted $K_{\mathrm{S}}$ estimates is taken as the consensus adjusted ortholog $K_{\mathrm{S}}$ estimate. The associated standard deviation is again computed via error propagation rules. Alternatively, *ksrates* can be configured to get the final rate-adjusted ortholog $K_{\mathrm{S}}$ estimate from the "best" outgroup species, thought of as most likely providing the most reliable ortholog $K_{\mathrm{S}}$ estimate decomposition. This "best" outgroup is taken to be the outgroup with the shortest O–C branch length (Supplementary Fig. 5), i.e. smallest $K_{\mathrm{OC}}$, which is internally computed using an equation similar to Equation 2. The more closely related the outgroup is to the two ingroup species and/or the lower its rate is, the better it is as an outgroup reference species. One should thus strive to pick such species to compile a good input dataset.

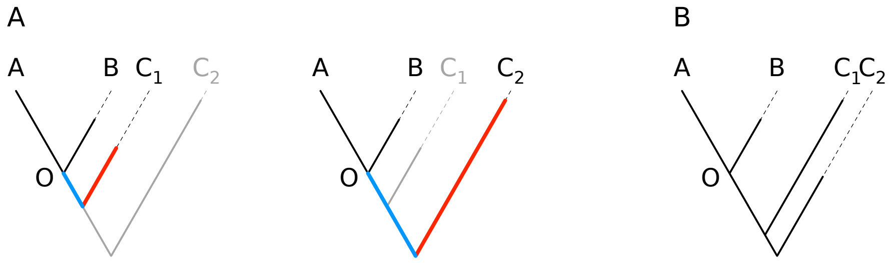

**Supplementary Figure 5.** Examples of outgroup choice. In the example tree in panel A, species $C_1$ is a better outgroup candidate than species $C_2$ because it is both more closely related to the two ingroup species A and B (shorter blue branch) and it has a lower synonymous substitution rate (shorter red branch), branch length O–$C_1$ < branch length O–$C_2$. In the example tree in panel B, species $C_2$ is the better outgroup candidate. Even though it is less closely related to the ingroup species pair A and B than species $C_1$, it does have a much lower synonymous substitution rate than species $C_1$, so that the combined effect (branch length O–$C_2$ < branch length O–$C_1$) makes species $C_2$ the better choice.

All calculated values for all species trios, including adjusted ortholog $K_S$, $K_{OA}$ and $K_{OC}$, are also written as tabular data to an output file (see Supplementary Table 1). This table can be inspected for consistency and e.g. used to spot outliers, which may suggest modifications to the analysis settings are necessary (see tool documentation at https://ksrates.readthedocs.io/).

| Node | Focal_Species | Sister_Species | Out_Species | Adjusted_Mode | Adjusted_Mode_SD | Original_Mode | Original_Mode_SD | Ks_Focal | Ks_Sister | Ks_Out |
|---|---|---|---|---|---|---|---|---|---|---|
| 2 | E. guineensis | O. sativa | A. officinalis | 0.659037 | 0.024384 | 1.531945 | 0.015001 | 0.329519 | 1.202427 | 0.795483 |
| 2 | E. guineensis | O. sativa | P. equestris | 0.654051 | 0.028293 | 1.531945 | 0.015001 | 0.327025 | 1.204920 | 1.071205 |
| 2 | E. guineensis | O. sativa | H. bullata | 0.652932 | 0.017413 | 1.531945 | 0.015001 | 0.326466 | 1.205479 | 0.444204 |
| 2 | E. guineensis | O. sativa | C. autumnale | 0.741159 | 0.020036 | 1.531945 | 0.015001 | 0.370579 | 1.161366 | 0.595879 |

**Supplementary Table 1.** Example rows from the output file with the ortholog $K_S$ rate-adjustment table obtained for focal species *Elaies guineensis*. Each row shows the $K_S$ adjustment data between *E. guineensis* and *Oryza sativa* from a species trio with a different outgroup (*Asparagus officinalis*, *Phalaenopsis equestris*, *Helonias bullata* or *Colchicum autumnale*). Column `Ks_Focal` contains the branch-specific $K_S$ contributions of the *E. guineensis* lineage, $K_{OA}$, and column `Ks_Sister` contains the branch-specific $K_S$ contributions of the *O. sativa* lineage, $K_{OB}$, to the original ortholog $K_S$ estimates, $K_{AB}$, in column `Original_Mode`. Column `Ks_Out` contains the contributions of the given outgroup lineage, $K_{OC}$, to $K_{AC}$ and $K_{BC}$, i.e. the $K_S$ length of the O–C branch for the given outgroup.

Ortholog $K_S$ adjustment results in a shift of the estimated $K_S$ position of a speciation event proportional to the estimated substitution rate difference between the diverged lineage and the lineage of the focal species. In case the focal species has a lower rate, the adjustment scales down the excess contribution of the diverged species to the $K_{AB}$ value, and the adjusted divergence $K_S$ estimate will be smaller than the original estimate. In case the focal species has a higher rate, the adjustment scales up the lower contribution of the diverged species to the $K_{AB}$ value, and the adjusted divergence $K_S$ estimate will be larger than the original estimate.

#### 2.3 Mixture modeling of paralog $K_S$ distributions

The interpretation of mixed paralog–ortholog $K_S$ distributions is sometimes challenged by the fact that paralog WGD peaks are often not clearly distinguishable due to progressive WGD signal erosion over time and due to potential overlaps between peaks of successive WGDs. In order to more objectively define the $K_S$ age of WGD peaks in the paralog $K_S$ distribution(s) of the focal species, *ksrates* uses various forms of mixture modeling. Depending on the available input data and analysis configuration (`paranome` and `collinearity` parameters in the *ksrates* configuration file), different methods are applied to analyze the paralog $K_S$ distribution(s) of the focal species (Supplementary Table 2).

If collinearity analysis is performed (`collinearity = yes`), the default method used is always a clustering approach based on the anchor-pair $K_S$ values in collinear segment pairs, which we call anchor $K_S$ clustering. Lognormal mixture modeling of the anchor-pair $K_S$ distribution and mixture modeling of the whole-paranome $K_S$ distribution (if `paranome = yes`) are optional. Otherwise, if only the whole-paranome analysis is performed (`collinearity = no`), exponential-lognormal mixture modeling of the whole-paranome $K_S$ distribution is used by default. Lognormal-only mixture modeling is optional.

Lognormal mixture modeling is not used as the default choice to model anchor-pair $K_S$ distributions because the alternative method (anchor $K_S$ clustering) applies the same mixture modeling algorithm on a biological unit, i.e. the segment pair (see below). For the whole-paranome $K_S$ distribution, exponential-lognormal mixture modeling is preferred over lognormal-only mixture modeling because the latter cannot adequately capture the roughly exponentially distributed small-scale duplication background (which is absent in anchor-pair $K_S$ distributions).

| Method | Collinearity-only | Paranome-only | Collinearity and paranome |
|---|---|---|---|
| Anchor $K_S$ clustering | ✓ | | ✓ |
| Exponential-lognormal mixture model | | ✓ | (✓) |
| Lognormal mixture model on anchor pairs | (✓) | | (✓) |
| Lognormal mixture model on paranome | | (✓) | (✓) |

**Supplementary Table 2.** Mixture modeling techniques used to detect potential WGD signals in paralog $K_S$ distributions. Default methods are marked by ✓, optional methods are marked by (✓). The applied methods depend on the configured analysis type (`collinearity` and/or `paranome`); optional methods can be turned on with an additional configuration setting.

##### 2.3.1 Anchor $K_S$ clustering

In order to classify anchor-pair $K_S$ values into groups tentatively representing different WGDs in the focal species' ancestry, *ksrates* uses a clustering approach. *ksrates* does not cluster the anchor-pair $K_S$ values directly, but instead clusters median $K_S$ values for the collinear segment pairs, i.e. pairs of sequence regions with conserved gene content and order, that the anchor pairs reside on. The collinear segments are detected and aligned by i-ADHoRe (Proost *et al.*, 2012), generating so-called multiplicons (Supplementary Fig. 6). The number of segments aligned in a multiplicon defines its level. A multiplicon may contain segment pairs that originated through different WGD events, but each pair of segments in a multiplicon traces back to a single WGD event.

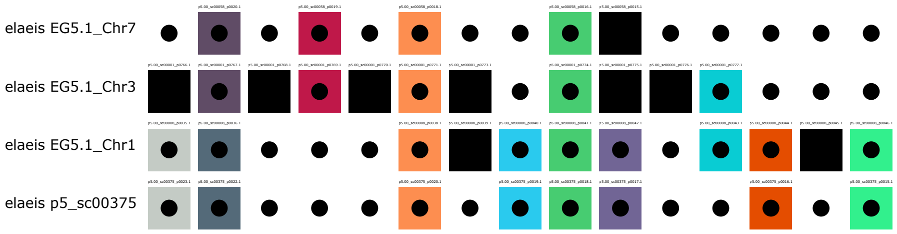

**Supplementary Figure 6.** Example of an i-ADHoRe multiplicon obtained for *Elaeis guineensis*, composed of four collinear segments (multiplicon level 4). Aligned anchor genes are depicted by boxes of the same color. The aligned anchor genes between any pair of collinear segments are called anchor pairs. Non-anchor genes in the segments are shown in black and alignment gaps in white.

Before the clustering of segment pairs, a cleaning step is performed. Some segment pairs are (partially) redundant because of the way multiplicons are generated. In the filtering step, segment pairs are compared between multiplicons and redundant ones are discarded according to the following criteria:

- Segment pairs whose anchor-pair list is a subset of the anchor-pair list of another segment pair are removed because they are fully redundant.
- If a segment pair has an anchor-pair list that partially overlaps ($> 1/3$) with the anchor-pair list of another segment pair, the segment pair with the shorter anchor-pair list is removed.

Smaller overlaps are tolerated in order not to lose too much collinear information. Note that the unitary element to be removed during the filtering is the entire segment pair and not individual redundant anchor pairs on it, to be consistent with the fact that collinearity traces are structured in blocks.

After obtaining the cleaned dataset, each segment pair is assigned a representative $K_S$ age. Even though all anchor pairs laying on a segment pair result from the same large-scale duplication, the $K_S$ estimates of the individual anchor pairs can vary substantially, e.g. due to stochastic effects in the synonymous substitution process, $K_S$ saturation effects and $K_S$ estimation errors (Vanneste *et al.*, 2013), and the resulting estimates frequently contain outliers. Anchor-pair $K_S$ lists with more than 5 elements are therefore pruned by removing the values falling outside the interval defined by the median $\pm$ median absolute deviation. Then, the representative $K_S$ age of each segment pair is computed as the median of the remaining anchor-pair $K_S$ list. The segment-pair median $K_S$ values are log-transformed and then clustered through Gaussian mixture modeling (GMM), which is equivalent to employing lognormal mixture modeling on the untransformed data. WGD peaks in $K_S$ distributions are preferentially modeled by lognormal distributions due to the fact that they generally exhibit positive skewness (Tiley *et al.*, 2018; Morrison, 2008). The clustered datapoints are back-transformed for cluster visualization on the original non-log $K_S$ scale (Supplementary Fig. 7A).

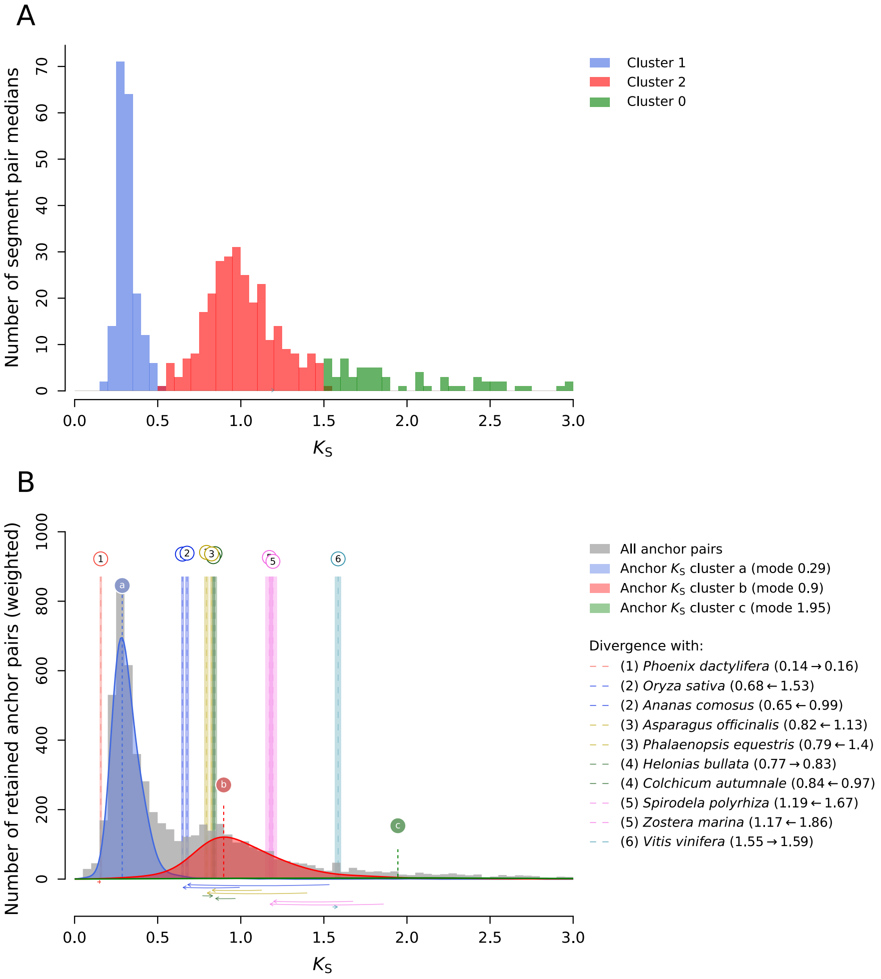

**Supplementary Figure 7.** Panel A shows a lognormal mixture model clustering (implemented as a Gaussian mixture model clustering on log-scale data) of collinear segment pair $K_S$ medians for *Elaeis guineensis*, where three clusters have been detected (blue, red and green). Panel B shows the mixed paralog–ortholog $K_S$ plot after each median has been substituted by its original anchor-pair $K_S$ list. The three $K_S$ clusters (filled blue, red and green KDE curves) are labeled with letters. The original anchor-pair $K_S$ distribution is visible as a gray histogram in the background. Substitution-rate-adjusted ortholog $K_S$ estimates of divergence events are visualized as in Fig. 1A in the main text.

GMM requires the number of clusters to be found to be set in advance. Usually, GMM is performed multiple times with a different target number of clusters, and the best number of clusters is selected based on, e.g. the Bayesian Information Criterion (BIC) or Akaike's Information Criterion (AIC). As this strategy in practice tends to overestimate the number of clusters (and hence WGDs), *ksrates* follows an alternative, more pragmatic approach. As the number of clusters should reflect the number of (detectable) WGDs in the lineage under study, *ksrates* estimates this number of WGDs beforehand based on the highest multiplicon level reached in the collinearity analysis. For example, the presence of 8 collinear segments can be explained by three whole-genome duplications. This approach is limited by the fact that a given maximum multiplicon level may be caused by different duplication scenarios, in particular taking into account that extensive post-WGD rearrangements and duplicate gene and segment losses are known to occur over evolutionary time. A maximum multiplicon level of 8 may for instance be caused by three whole-genome duplications or by two whole-genome triplications (theoretical multiplicon level 9) followed by extensive gene loss. *ksrates* takes an upper boundary of the minimum number of duplication events across these scenarios as the GMM cluster number estimate, by assuming only whole-genome duplications have happened. A maximum multiplicon level of 9 for instance requires at least 4 whole-genome duplications, although it may be explained by, e.g. 2 whole-genome triplications or 2 whole-genome duplications and a triplication. Superfluous clusters caused by overestimating the number of WGDs are filtered out afterwards to the extent possible, as described below.

Given the cluster number estimated from the maximum multiplicon level, the GMM is initialized and fitted on the segment-pair medians $K_S$ dataset multiple times, and the best model is taken to be the one with the largest log-likelihood. Subsequently, the segment-pair median $K_S$ values are replaced by the original $K_S$ list for the segment pair to obtain the anchor $K_S$ clusters (Supplementary Fig. 7B). After initial clustering, anchor $K_S$ clusters for which a link to a real WGD event is ambiguous or unlikely are removed from the dataset, along with the segment pairs defining them. Removed clusters meet at least one of the following empirical criteria (thresholds are based on cluster shapes obtained for test species, see Section 4):

- The cluster is poorly populated (it contains <= 10% of the total number of $K_S$ values). These are likely clustering artifacts, in particular when the cluster has a young median $K_S$ age.
- The cluster $K_S$ estimate is too old to be reliably associated with a WGD (the median $K_S$ of the cluster is >= 3). Although ancient WGD events may have happened beyond $K_S$>= 3, finding reliable evidence for those in $K_S$ plots is challenging, as $K_S$ peaks beyond $K_S$>= 3 are increasingly likely to be artifacts caused by $K_S$ saturation effects (Vanneste *et al.*, 2013).
- The cluster has a flat, stretched-out peak signal (the inter-quartile range of the cluster is >= 1.1 $K_S$). These are likely artifacts, in particular when the cluster has a young median $K_S$ age.

These criteria mostly remove clustering artifacts in the anchor $K_S$ distribution tail and small $K_S$ clusters that overfit particular distribution features, often at $K_S$ values close to 0 or in between $K_S$ peaks that represent genuine WGDs (compare Supplementary Fig. 7B with Supplementary Fig. 13). If one or more clusters are filtered out, a second round of GMM is performed on the remaining segment pairs using the remaining number of clusters. For example, the green 'c' cluster in Supplementary Fig. 7 is removed because it matches two of the aforementioned criteria (see also Fig. 1A in the main text).

###### 2.3.2 Exponential-lognormal mixture model

*ksrates* implements the expectation-maximization (EM) algorithm described in Zhang *et al.* (2019) to fit exponential-lognormal mixture models to whole-paranome $K_S$ distributions. The lognormal components model putative WGD peaks, and their modes are used as proxy for the WGD age (Supplementary Fig. 8). The exponential component models the L-shaped background $K_S$ distribution generated by small-scale duplications (Blanc and Wolfe, 2004) (compare Supplementary

Fig. 8 with Supplementary Fig. 12).

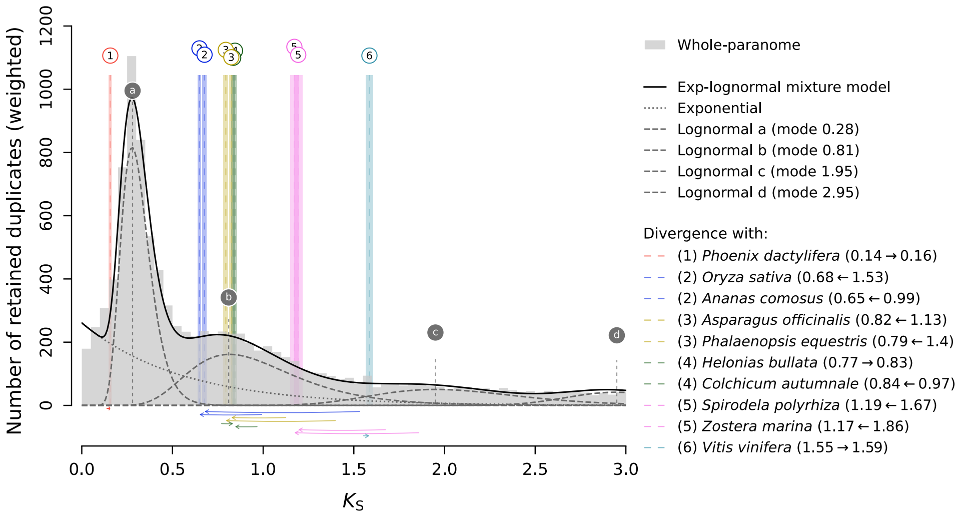

**Supplementary Figure 8.** Mixed paralog–ortholog $K_S$ plot showing the whole-paranome $K_S$ distribution for *Elaeis guineensis* in light gray with superimposed exponential-lognormal mixture model. The overall mixture model (dark solid line) is composed of an exponential component (dotted gray curve) and lognormal components (dashed gray curves). Each lognormal component is labeled with a letter (vertical dashed gray lines with circular labels). Components a and b are matching the two WGD peaks visible in the paralog $K_S$ distribution. The "buffer" component d covers the right-most side of the $K_S$ range. Substitution-rate-adjusted ortholog $K_S$ estimates of divergence events are visualized as in Fig. 1A in the main text. Note that the exponential-lognormal model positions event b around the time the *E. guineensis* lineage diverged from Asparagales (*P. equestris* and *A. officinalis*). The anchor-pair $K_S$ clustering shown in Fig. 1A instead positions the b event ($\tau$ WGD) at the commonly accepted phylogenetic position during the early diversification of monocots.

Duplication events are often not represented by a single $K_S$ value in the whole-paranome $K_S$ distribution, but by a number of $K_S$ values with fractional weights summing to 1 (see Section 2.1 and Maere *et al.* (2005)). As the EM algorithm used for mixture model optimization cannot handle such fractional weights, mixture modeling is not run on the original weighted paralog $K_S$ distribution, but on a deconvoluted version. To this end, the weighted paralog $K_S$ values in each bin of width 0.01 in the original distribution are replaced by $n$ $K_S$ values equal to the bin central value, where $n$ is the sum of weights of the Ks values in the bin rounded to the nearest integer. To avoid edge effects caused by the truncation of the distribution's right tail, the $K_S$ range in which model fitting is applied (customizable, default $K_S$ range = [0,5]) is internally increased by 0.5 $K_S$. Moreover, an extra "buffer" lognormal component is initialized at the right boundary of this fitting range to prevent that other components stretch towards higher values in an attempt to model the hard-to-fit right tail.

Since adequate initialization of the component parameters is crucial for obtaining decent mixture modeling results, *ksrates* uses three different initialization approaches and ultimately chooses the best one based on BIC:

**Data-driven initialization** In this method the initialization of component parameters is guided by the shape of the whole-paranome $K_S$ distribution. The initialization of the exponential component takes advantage of the fact that the intercept of the exponential probability density function with the y-axis is equal to the decay rate of the function (Supplementary Fig. 9). This intercept is approximated by the height of the first bin in a probability density histogram of the weighted whole-paranome $K_S$ distribution with bin width 0.1 $K_S$, which is narrow enough to have a good resolution at the left edge of the $K_S$ distribution but at the same time wide enough to avoid potential biases caused by an artificially high density close to $K_S = 0$ followed by a sudden drop (often caused by sequencing artifacts).

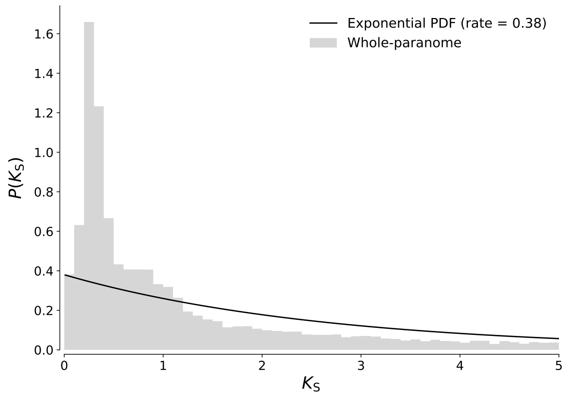

**Supplementary Figure 9.** The decay rate of the exponential probability density function (black curve) is equal to the intercept with the y-axis. The light gray histogram is the whole-paranome $K_S$ probability density histogram for *Elaeis guineensis* with a bin width of 0.1 $K_S$.

To initialize the lognormal means and standard deviations, the paranome $K_S$ distribution is first log-transformed so that the putative lognormal WGD peaks assume a Gaussian shape. Then, the transformed distribution is searched for peaks as follows. First, a KDE (with Gaussian kernel and bandwidth at 40% of the bandwidth determined by Scott's rule (Scott, 1992)) is computed on an extended distribution in which data on the original right boundary is reflected to the other side, to prevent KDE edge effects (Supplementary Fig. 10). Subsequently, a smoothing spline is computed on the KDE in order to smooth out small irregularities (Supplementary Fig. 10).

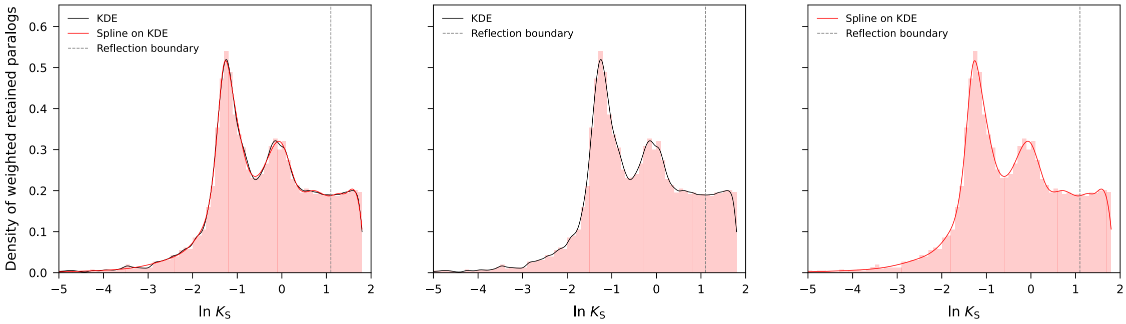

**Supplementary Figure 10.** KDE (black line) and spline (red line) obtained from the log-transformed whole-paranome $K_S$ distribution of *Elaeis guineensis* (light red histogram). The spline smoothes out irregularities in the KDE and is used for data-driven peak detection. The underlying paralog distribution has been partially reflected across the right boundary (dashed vertical line) to account for edge effects.

This spline may still exhibit small noise peaks (Supplementary Fig. 11A). In an attempt to filter away these small peaks and retain only presumedly real WGD peaks, the distribution is mirrored around each peak in both directions (Supplementary Fig. 11B and C). Then, the prominence of each mirrored peak is measured, i.e. how much the signal stands out from the surrounding baseline. The peak is retained as a possible WGD peak only if its prominence is significant in at least one reflection, with a significance threshold set to 0.06 on the basis of empirical results coming from test species (see Section 4). The ln $K_S$-coordinates of significant peaks are then taken as the mean of Gaussian components. The component standard deviation is initialized as the peak half width, calculated at 60% of the prominence height in the reflection with the highest prominence (Supplementary Fig. 11B and C) (chosen on the basis of empirical results coming from test species, see Section 4).

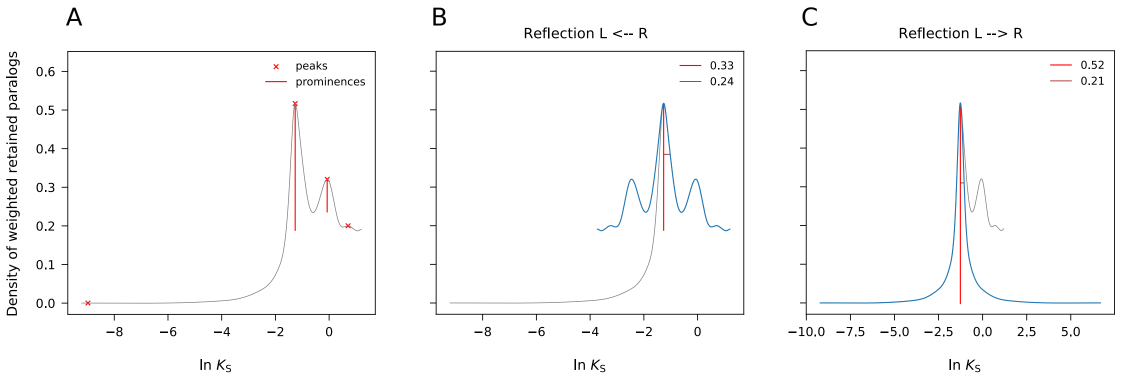

**Supplementary Figure 11.** Panel A shows a whole-paranome smoothing spline (gray curve) for *Elaeis guineensis* with peaks (red crosses) and prominence heights (red lines) indicated. Panels B and C show the reflections (blue curves) around the tallest peak in both directions. The peak has a significant prominence height (value in legend) in both reflections and its ln $K_S$ coordinate is therefore used as a normal component mean. The peak half width at 60% prominence height (horizontal dark red line, value in legend) in the reflection with the highest prominence is used to initialize the standard deviation of the component.

In case of overlapping WGD signals, it is possible that the computed peak width stretches across multiple signals, overestimating the width of the putative signal underlying the reflected peak. To avoid this pitfall, the component standard deviation is capped at 0.75 (ln $K_S$ units). The calculated Gaussian component means and standard deviations are then used to initialize the corresponding lognormal components in the original $K_S$ space. The EM algorithm finally proceeds with fitting all initialized components in the original $K_S$ space.

**Random initialization** In the random initialization method, the components are initialized by randomly drawing their parameter values from a range of reasonable values empirically obtained from test species (see Section 4):

- the exponential rate is chosen between 0.2 and 1 in steps of 0.1
- the normal mean is chosen between −0.5 and 0.9 in steps of 0.1
- the normal standard deviation is chosen between 0.3 and 0.9 in steps of 0.1

A mixture model is by default fitted with two to five random components (including the "buffer" component which is always present, customizable range). For each number of components, the mixture model is initialized multiple times (customizable parameter) and the best fit is chosen according to the lowest BIC score.

**Hybrid initialization** For the hybrid method, the mixture model is initialized with the components previously computed through the data-driven approach, but with the addition of a single lognormal component whose parameters are randomly drawn from the ranges used in the random initialization method. This method is an attempt to reintroduce peaks that were overlooked by the data-driven approach. The mixture model is initialized multiple times (customizable parameter) and the best fit is chosen according to the lowest BIC score.

**Model evaluation** After having run models with all three initialization approaches, the model with the lowest BIC value is incorporated in the mixed paralog–ortholog $K_S$ distribution plot (Supplementary Fig. 8).

##### 2.3.3 Lognormal mixture models

The lognormal mixture model is optionally performed for both the whole-paranome $K_S$ distribution (Supplementary Fig. 12) and the anchor pair $K_S$ distribution (Supplementary Fig. 13) of the focal species. A Gaussian mixture model is fitted on deconvoluted log-transformed $K_S$ data and subsequently converted to the correspondent lognormal mixture model in the original $K_S$ space. As in the exponential-lognormal mixture model (see previous section), the $K_S$ fitting range (customizable) is internally increased by 0.5 $K_S$ to avoid edge effects caused by the truncation of the distribution's right tail. The mixture models are by default fitted with two to five components (customizable range). For each number of components, the mixture model is initialized multiple times (customizable parameter) and the best fit is chosen according to the largest log-likelihood. Among the resulting models (one for each number of components), the best fitting model is taken to be the one with the lowest BIC score.

Lognormal mixture modeling is only applied optionally due to its tendency to over-estimate the number of components. This is particularly (but not exclusively) the case when modeling whole-paranome $K_S$ distributions, which contain a small-scale duplication background that cannot be captured adequately by lognormals (compare Supplementary Fig. 12 with Supplementary Fig. 8).

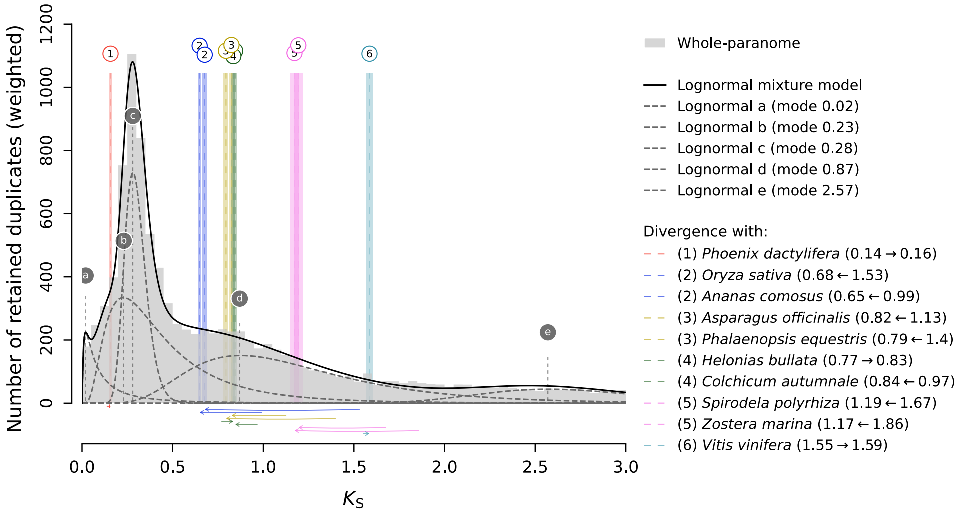

**Supplementary Figure 12.** Mixed paralog–ortholog $K_S$ plot showing the whole-paranome $K_S$ distribution for *Elaeis guineensis* in light gray superimposed with the best-fitting lognormal-only mixture model. The overall lognormal mixture model (solid black curve) is composed of multiple lognormal components (dashed gray curves), which are labeled with letters (vertical dashed gray lines with circular labels). Substitution-rate-adjusted ortholog $K_S$ estimates of divergence events are visualized as in Fig. 1A in the main text.

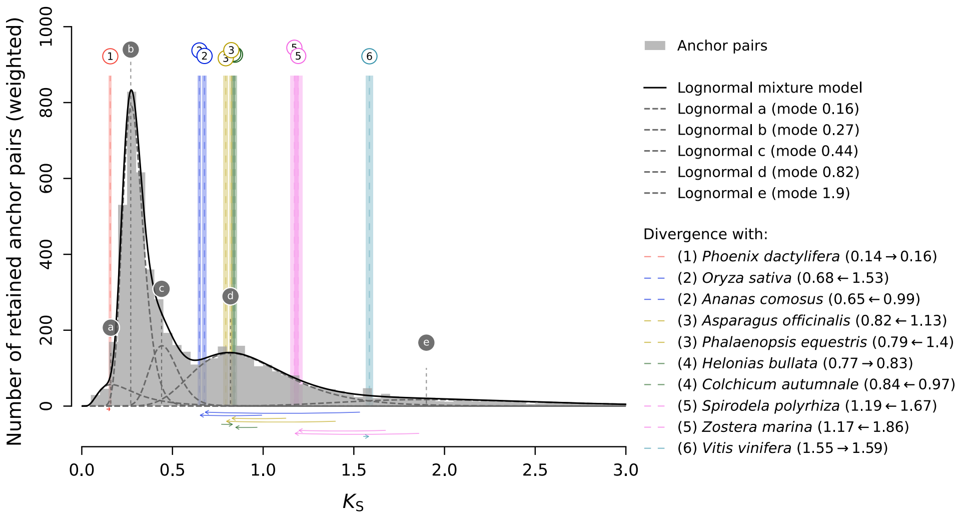

**Supplementary Figure 13.** Mixed paralog–ortholog $K_S$ plot showing the anchor pair $K_S$ distribution for *Elaeis guineensis* in dark gray superimposed with the best-fitting lognormal-only mixture model. The overall lognormal mixture model (solid black curve) is composed of multiple lognormal components (dashed gray curves), which are labeled with letters (vertical dashed gray lines with circular labels). Substitution-rate-adjusted ortholog $K_S$ estimates of divergence events are visualized as in Fig. 1A in the main text.

### 3 Data sources

Sources of input CDS FASTA and GFF data for all species used in the use case analysis (Fig. 1 in the main text and most of the Supplementary Figures here) are provided in Supplementary Table 3. Genome data were used for most of the species, while transcriptome data were used for *Helonias bullata*, *Colchicum autumnale* and *Illicium floridanum*.

### 4 Parameters used

For the *ksrates* use case analysis on *Elaeis guineensis* presented in the main text and this supplement we used the following parameter settings. The complete *ksrates* configuration files have been deposited together with all data underlying this article in Zenodo, at https://dx.doi.org/10.5281/zenodo.4717851.

**Ortholog $K_S$ distributions and rate adjustment** The Newick tree (((((((*elaeis*, *phoenix*), (*oryza*, *ananas*)), (*asparagus*, *phalaenopsis*)), (*helonias*, *colchicum*)), (*spirodela*, *zostera*)), *vitis*), *illicium*) was used as the input phylogeny (parameter `newick_tree`, using short species IDs), with *E. guineensis* (*elaeis*) as the focal species (parameter `focal_species`). Ortholog $K_S$ distributions were constructed up to the default value of $K_S$=10 (parameter `max_ks_orthologs`) and plotted up to the default value of $K_S$=5 (parameter `x_axis_max_limit_orthologs_plots`) using the default histogram bin width of 0.1 (parameter `bin_width_orthologs`). Modes of the ortholog $K_S$ distributions were estimated using 200 bootstrapped KDEs (default value for parameter `num_bootstrap_iterations`). The KDE bandwidth for each bootstrapped dataset was first selected using Scott's rule (Scott, 1992) and then multiplied by a random number between 0.5 and 0.9 since the bandwidth selected by Scott's rule resulted in oversmoothed ortholog $K_S$ distributions. Up to four closest outgroup species (default value for parameter `max_number_outgroups`) were used to calculate multiple rate-adjusted ortholog $K_S$ estimates per divergence event, and the mean (default value for parameter `consensus_mode_for_multiple_outgroups`) of these multiple adjusted $K_S$ estimates was used as consensus. Plotting of horizontal arrows below the mixed paralog–ortholog

$K_S$ plots was turned on (expert parameter `plot_adjustment_arrows`) to visualize the shifts of the ortholog $K_S$ estimates resulting from the rate-adjustment.

| Species | Version / Accession | Source database |
|---|---|---|
| *Elaeis guineensis* | EG5.1 | Monocots PLAZA 4.5 (Locus FASTA Data and Structural Annotation GFF) https://bioinformatics.psb.ugent.be/plaza/versions/plaza_v4_5_monocots/download |
| *Phoenix dactylifera* | GCF_000413155.1 | NCBI ftp://ftp.ncbi.nlm.nih.gov/genomes/all/GCF/000/413/155/GCF_000413155.1_DPV01 |
| *Oryza sativa ssp. japonica* | v7_JGI | Monocots PLAZA 4.5 (Locus FASTA Data) https://bioinformatics.psb.ugent.be/plaza/versions/plaza_v4_5_monocots/download |
| *Ananas comosus* | v3 | Monocots PLAZA 4.5 (Locus FASTA Data) https://bioinformatics.psb.ugent.be/plaza/versions/plaza_v4_5_monocots/download |
| *Asparagus officinalis* | v1.1 | Monocots PLAZA 4.5 (Locus FASTA Data) https://bioinformatics.psb.ugent.be/plaza/versions/plaza_v4_5_monocots/download |
| *Phalaenopsis equestris* | v1.0 | Monocots PLAZA 4.5 (Locus FASTA Data) https://bioinformatics.psb.ugent.be/plaza/versions/plaza_v4_5_monocots/download |
| *Helonias bullata* | OOSO | One Thousand Plants Project (1KP) http://www.onekp.com/public_data.html |
| *Colchicum autumnale* | QNPH | One Thousand Plants Project (1KP) http://www.onekp.com/public_data.html |
| *Spirodela polyrhiza* | v2 | Monocots PLAZA 4.5 (Locus FASTA Data) https://bioinformatics.psb.ugent.be/plaza/versions/plaza_v4_5_monocots/download |
| *Zostera marina* | v2.2 | Monocots PLAZA 4.5 (Locus FASTA Data) https://bioinformatics.psb.ugent.be/plaza/versions/plaza_v4_5_monocots/download |
| *Vitis vinifera* | Genoscope.12X | Monocots PLAZA 4.5 (Locus FASTA Data) https://bioinformatics.psb.ugent.be/plaza/versions/plaza_v4_5_monocots/download |
| *Illicium floridanum* | VZCI | One Thousand Plants Project (1KP) http://www.onekp.com/public_data.html |
| *Musa acuminata* | Banana Genome v2.0 | Monocots PLAZA 4.5 (Locus FASTA Data) https://bioinformatics.psb.ugent.be/plaza/versions/plaza_v4_5_monocots/download |
| *Arabidopsis thaliana* | Araport11 | Dicots PLAZA 4.5 (Locus FASTA Data) https://bioinformatics.psb.ugent.be/plaza/versions/plaza_v4_5_dicots/download |

**Supplementary Table 3.** Data source table. The left column specifies the species, the middle column specifies the genome version or the data accession code and the right column specifies the source database and the link to the associated website's download page.

**Paralog $K_S$ distributions** The construction of both whole-paranome and anchor-pair $K_S$ distributions was turned on (parameters `paranome` and `collinearity`) for the focal species *E. guineensis*. The built-in default settings of the *wgd* package for paralog and ortholog detection and $K_S$-value estimation and for i-ADHoRe execution were used except that paralog gene families with more than 200 members were excluded (200 is the default value of the *ksrates* expert parameter `max_gene_family_size`, which is higher than the default setting in *wgd*). This resulted in the exclusion of four paralog gene families with a size larger than 200. To extract the relevant genome structural information from the *E. guineensis* GFF file the keywords "mRNA" (parameter `gff_feature`) and "ID" (parameter `gff_attribute`) were used. *E. guineensis* paralog $K_S$ distributions were constructed up to the default value of $K_S$=5 (parameter `max_ks_paralogs`), but plotted only up to the value of $K_S$=3 (parameter `x_axis_max_limit_paralogs_plots`) using a histogram bin width of 0.05 (parameter `bin_width_paralogs`). The bandwidth for the KDEs of the whole-paranome and anchor-pair $K_S$ distributions (see Supplementary Fig. 2) was first selected using Scott's rule and the selected value was then multiplied by 0.3 (expert parameter `kde_bandwidth_modifier`) to obtain a tighter fit of the distributions.

**Paralog mixture modeling** All mixture modeling methods provided by *ksrates*, including the optional ones, were turned on (expert parameter `extra_paralogs_analyses_methods`). Default parameter settings were used for the EM algorithm: 10 initializations (expert parameter `num_mixture_model_initializations`), a maximum of 300 EM iterations (expert parameter `max_mixture_model_iterations`), a maximum of five lognormal components (expert parameter `max_mixture_model_components`), and the paralog $K_S$ distributions were fit up to a maximum value of $K_S$=3 (expert parameter `max_mixture_model_ks`). The convergence value for the EM algorithm is $1 \times 10^{-6}$.

The *ksrates* mixture modeling methods have been finetuned on paralog $K_S$ distributions generated for the following set of test species next to *E. guineensis*: *Arabidopsis thaliana*, *Oryza sativa*, *Musa acuminata*, *Ananas comosus* and *Asparagus officinalis* (data not shown).

### 5 Software availability and requirements

*ksrates* is open-source software implemented in Python 3 and as a Nextflow pipeline, and is available for free under the GNU GPL v3 license from the public GitHub repository https://github.com/VIB-PSB/ksrates. We provide *ksrates* as Singularity and Docker containers that bundle all required external software dependencies (except Nextflow) and as a simple Python package. In principle, *ksrates* runs on any Linux or macOS system or on Windows with Windows Subsystem for Linux 2 (WSL2) or a virtual machine installed. However, constructing multiple $K_S$ distributions is computationally demanding, and therefore we recommend the use of a computer cluster (or cloud platform) for any input phylogeny larger than the minimal 3-species phylogeny. The containers require Singularity or Docker software to be installed. Usage of the Nextflow pipeline requires Nextflow itself, Java 8 (or later) and Bash 3.2 (or later) to be installed. Usage of the Python package instead of one of the containers requires Python 3 and various Python packages to be installed, as well as external software dependencies of the *wgd* package (Zwaenepoel *et al.*, 2019) (as listed in Section 2.1). Detailed installation and usage instructions are included in the *ksrates* documentation. The documentation, a tutorial and example datasets are available via https://ksrates.readthedocs.io and https://github.com/VIB-PSB/ksrates.

The *ksrates* Nextflow pipeline allows to run the complete analysis automatically. Nextflow also makes it easy to configure the execution of the pipeline on a variety of computer clusters, such as Sun Grid Engine (SGE) or compatible clusters or clusters using the PBS/Torque family of batch schedulers, and cloud platforms, such as Amazon Web Services (AWS) or Kubernetes, although we did not test the latter. Alternatively, the individual steps of a *ksrates* analysis can be easily executed manually using a command-line interface available via the Singularity and Docker containers and the Python package. This provides more control over and customization of the analysis and allows the integration into existing genomics toolsets and workflows.

To get familiar with the use of *ksrates*, we provide two example datasets. A small test dataset composed of truncated sequence files can be used to perform a quick check whether *ksrates* is correctly installed and fully functional, which should take only a few minutes to run. The full example dataset contains the sequence data and template configuration files needed to reproduce the use case scenario described in the tutorial and documentation. This dataset is best run on a computer cluster, as processing it on an average laptop or desktop computer may take several hours.
